# Fully efficient, two-stage analysis of multi-environment trials with directional dominance and multi-trait genomic selection

**DOI:** 10.1101/2022.09.28.509884

**Authors:** Jeffrey B. Endelman

## Abstract

Plant breeders interested in genomic selection often face challenges to fully utilizing the multi-trait, multi-environment datasets they rely on for selection. R package StageWise was developed to go beyond the capabilities of most specialized software for genomic prediction, without requiring the programming skills needed for more general-purpose software for mixed models. As the name suggests, one of the core features is a fully efficient, two-stage analysis for multiple environments, in which the full variance-covariance matrix of the Stage 1 genotype means is used in Stage 2. Another feature is directional dominance, including for polyploids, to account for inbreeding depression in outbred crops. StageWise enables selection with multi-trait indices, including restricted indices with one or more traits constrained to have zero response. For a potato dataset with 943 genotypes evaluated over 6 years, including the Stage 1 errors in Stage 2 reduced the Akaike Information Criterion (AIC) by 29, 67, and 104 for maturity, yield, and fry color, respectively. The proportion of variation explained by heterosis was largest for yield but still only 0.03, likely because of limited variation for the genomic inbreeding coefficient. Due to the large additive genetic correlation (0.57) between yield and maturity, naïve selection on an index combining yield and fry color led to an undesirable response for later maturity. The restricted index coefficients to maximize genetic merit without delaying maturity were identified. The software and three vignettes are available at https://github.com/jendelman/StageWise.

## INTRODUCTION

During the first decade of the 21^st^ century, the focus of genomic selection research was the development of theory and methods (e.g., Meuwissen et al. 2001; Habier et al. 2007; Daetwyler et al. 2008; Bernardo and Yu, 2007; VanRaden 2008), and most researchers worked in animal rather than plant breeding. This changed in the following decade with the development of specialized software for genomic prediction, including rrBLUP (Endelman 2011), GAPIT (Lipka et al. 2012), synbreed (Wimmer et al. 2012), BGLR (Pérez and de los Campos 2014), and sommer (Covarrubias-Pazaran 2016). Collectively, these software publications have been cited several thousand times, which reflects their enabling role for the adoption of genomic selection, particularly in plant breeding.

However, these packages were not designed to handle the full complexity of plant breeding data, spanning multiple environments with different experimental designs, heritabilities, and spatial models for non-genetic variation. The challenge of properly analyzing multi-environment datasets existed before genomic selection, which led to the concept of a two-stage analysis (Frensham et al. 1997). In Stage 1, genotype means are estimated as fixed effects for each environment, which become the response variable in Stage 2. The errors of the Stage 1 estimates are typically different, and failure to account for this in Stage 2 leads to sub-optimal results (Möhring and Piepho 2009). Nonetheless, Stage 1 errors are commonly ignored in published studies; I believe a major reason is the additional programming skill required to follow best practice with the available software.

The goal of the current research was to develop genomic selection software with a relatively simple interface that still allows for a fully efficient, two-stage analysis. The description “fully efficient” implies the full variance-covariance matrix of the Stage 1 genotype means is used in Stage 2, not just a diagonal approximation (Piepho et al. 2012; Damesa et al. 2017). The result is StageWise, a package for the R computing environment (R Core Team 2022) that works for any ploidy and incorporates advanced features, such as directional dominance and multi-trait selection indices.

## METHODS

### Single trait with homogeneous GxE

The response variable for Stage 2 is the Stage 1 BLUEs for the effect of genotype in environment. The mixed model with homogeneous GxE can be written as

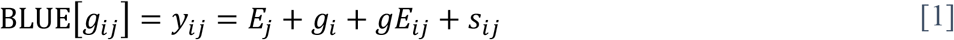

where *g*_*ij*_ is the genotypic value for individual (or clone) *i* in environment *j, E*_*j*_ is the fixed effect for environment *j, g*_*i*_ is the random effect for individual *i* across environments, and the GxE effect, *gE*_*ij*_, is actually the model residual (Damesa et al. 2017). The *s*_*ij*_ effect, which represents the Stage 1 estimation error, is multivariate normal with no free variance parameters: the variance-covariance matrix is the direct sum of the variance-covariance matrices of the Stage 1 BLUEs (Damesa et al. 2017). The *gE*_*ij*_ are independent and identically distributed (i.i.d.), which implies a single genetic correlation between all environments. Without marker data, the software assumes the *g*_*i*_ effects are i.i.d.

When marker data are provided, the software decomposes *g*_*i*_ into additive and non-additive values. The vector of additive values is multivariate normal with covariance proportional to a genomic additive matrix **G** (VanRaden 2008 Method 1, extended to arbitrary ploidy). If **W** represents the centered matrix of allele dosages (*n* individuals x *m* bi-allelic markers with frequencies *p* = 1–*q*), then for ploidy *ϕ*,

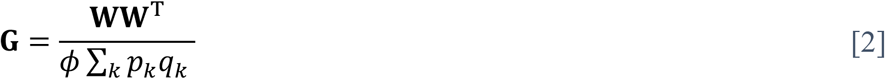

If a three-column pedigree is provided, **G** can be blended with the pedigree relationship matrix **A** (calculated using R package AGHmatrix [Amadeu et al. 2016]) to produce **H** = (1 − ω)**G** + ω**A**, for 0 ≤ ω ≤ 1 (Legarra et al. 2009; Christensen and Lund 2010). In addition to the additive polygenic effect, the user can indicate some markers should be included as additive (fixed effect) covariates in Eq. 1, to capture large effect QTL.

### Directional dominance

Two models for the non-additive genetic values are available. In the genetic residual model, the non-additive values are i.i.d. The other option is a directional (digenic) dominance model, which follows the classical framework of Fisher (1941) and Kempthorne (1957) and is a refinement of recent research (Vitezica et al. 2013; Xiang et al. 2016; Endelman et al. 2018; Batista et al. 2021). For a locus with two alleles designated 0/1, there are three digenic dominance effects *β*_00_, *β*_01_, *β*_11_, which equal the dominance deviation in diploids, but more generally for any ploidy are the coefficients for regressing the dominance deviation on diplotype dosage. (Higher order dominance effects for polyploids are not considered.) These dominance effects can be expressed in terms of a parameter that has no established name but may be called a digenic substitution effect, *β*, by analogy with the allele substitution effect *α* for additive effects. The *β* parameter represents the average change in dominance deviation per unit increase in dosage of the heterozygous diplotype:

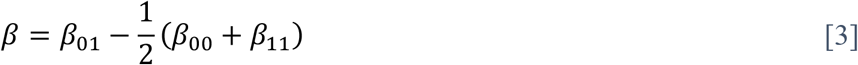

(This differs from the scaling in Endelman et al. [2018] by –2 so that *β* in Eq. 3 equals *d* in the classical diploid model of Vitezica et al. [2013].) Designating the frequency of allele 1 as *p* = 1 − *q*, the dominance effects can be expressed in terms of the substitution effect:

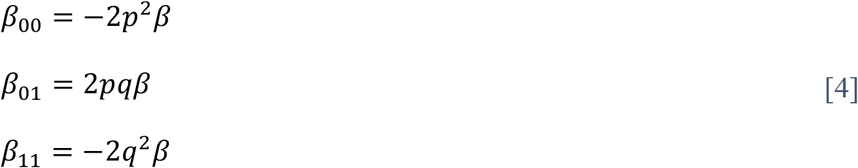

The dominance value of an individual is the sum of its dominance effects and can be written as *Qβ*, where the dominance coefficient *Q* for ploidy *ϕ* and allele dosage *X* (of allele 1) is

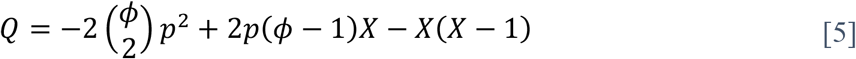

In Eq. 5, 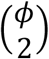 is the binomial coefficient. The dominance genetic variance, *V*_*D*_, is 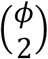 times the variance of the dominance effects, 4*p*^2^*q*^2^*β*^2^. Extending this framework to *m* loci, the dominance value is 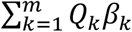, and the dominance variance is

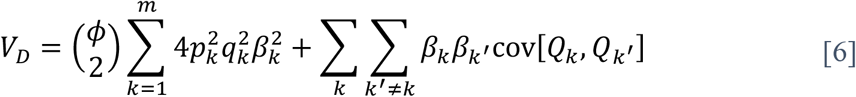

The first term in Eq. 6 is the dominance *genic* variance, which depends on allele frequencies but not LD between loci. The second term is the disequilibrium covariance, which can be positive or negative.

In classical quantitative genetics, the substitution effects are fixed parameters, but to compute dominance values by BLUP, we switch to viewing them as random normal effects (de los Campos et al. 2015), with mean *μ*_*β*_ and variance 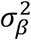. For a trait with no heterosis, *μ*_*β*_ = 0 (Varona et al. 2018). Let **Q** denote the *n* x *m* matrix of dominance coefficients for *n* individuals at *m* loci. The vector of dominance values **Qβ** is multivariate normal, with mean **Q1***μ*_*β*_ and variance-covariance matrix 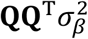. Equivalently, the dominance values can be written as

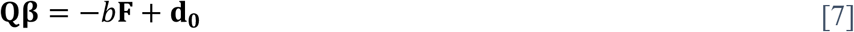

where **F** is a vector of genomic inbreeding coefficients, with regression coefficient *b* (positive value implies heterosis), and 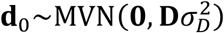 represents dominance in the absence of heterosis. The genomic dominance matrix **D** is defined by interpreting its variance component 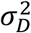 as the expected value of the classical dominance variance with respect to the substitution effects, assuming no overall heterosis. From Eq. 6 the result is

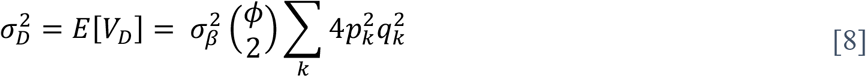

which leads to

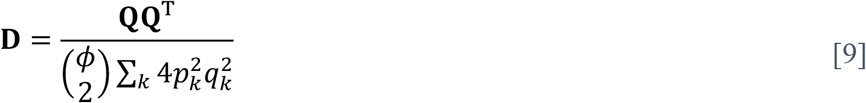

From Eq. 7, the vector of genomic inbreeding coefficients **F** is proportional to the row sum of **Q**. The correct scaling is derived by considering the expected value of *Q* (Eq. 5) in the classical sense (where genotypes are random and parameters are fixed), for a completely inbred population in which homozygotes of allele 1 occur with frequency *p*. Under these conditions, *E*[*X*] = *ϕp* and *E*[*X*^2^] = *ϕ*^2^*p*, which leads to 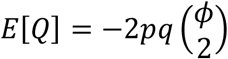. Extending this to multiple loci and equating the result to *F* = 1 sets the proportionality constant and leads to the following definition:

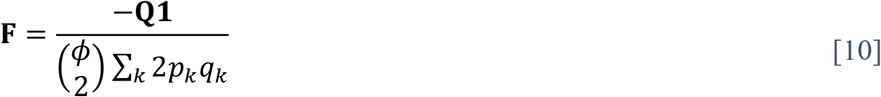

The vector of genomic inbreeding coefficients is included as a fixed effect covariate in the Stage 2 model. Inbreeding coefficients can also be computed from the diagonal elements of the additive relationship matrix (either A or G) according to (*G* − 1)/(*ϕ* − 1)(Henderson 1976; Gallais 2003; Endelman and Jannink 2012).

### Extension to multiple locations or traits

StageWise has the option of including a random effect *g*(*L*) in Stage 2 for genotype within location (or *L* can represent some other factor, such as management). Using the subscript *k* to designate location, the linear model (Eq. 1) becomes

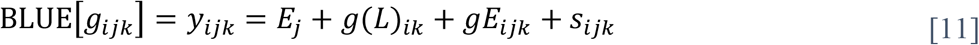

The *g*(*L*)_*ik*_ effect is modeled using a separable covariance structure, **I⨂Γ** in the absence of marker data, where the genetic covariance between locations **Γ** follows a second order factor-analytic (FA2) model. The FA2 model provides a good balance between statistical parsimony and complexity for many plant breeding applications, and *Stage2* returns the rotated and scaled factor loadings (Cullis et al. 2010). A heterogeneous variance model is used for *gE*_*ijk*_ (which is the model residual as before), with different variance parameters for each location.

When marker data are provided, genotypic value is partitioned into additive and non-additive values, and the FA2 model is still used for the additive covariance between locations. Attempts to use an FA2 model for non-additive values were unsuccessful in several datasets, and even with a compound symmetry model, the correlation parameter was always on the boundary (equal to 1). The non-additive correlation parameter was therefore fixed at 1 and accepted as a model limitation. When markers are included as fixed effect covariates, different regression coefficients are estimated for each location. Similarly, different regression coefficients for genomic inbreeding are estimated per location.

A similar framework is used for multi-trait analysis, with trait replacing location in Eq. 11, except that all trait covariance matrices are unstructured. In Stage 1, a separable covariance model is used for the residuals, and in Stage 2, the fixed effects for environment are trait-specific. When markers are used to partition additive and non-additive genetic value, separate unstructured covariance matrices are estimated for each. Multi-trait models are limited to the homogeneous GxE structure described for single trait analysis (i.e., the genetic correlation between all environments is the same, regardless of location).

### Proportion of variance explained

The aim is to quantify the proportion of variance (PVE) explained by each effect in the Stage 2 model, excluding the main effect *E*_*j*_ (which mirrors how heritability is calculated). The core idea is to compute variances based on the method of Legarra (2016), and the PVE is the variance of each effect divided by the sum. This is not a true partitioning of variance because the Stage 2 effects are not necessarily orthogonal.

First consider effects such as *gE*_*ij*_ and *s*_*ij*_ (Eq. 1), which are indexed by both genotype *i* and environment *j*. Representing these effects by vector **y** of length *t*, the variance is

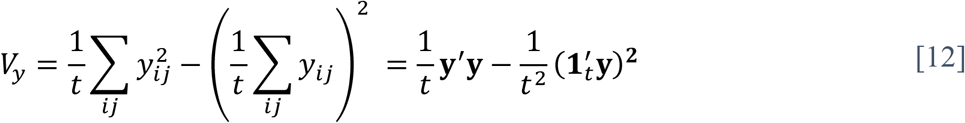

The symbol **1**_*t*_ in Eq. 12 is a *t* x 1 vector of 1’s. For multivariate normal (MVN) **y** with mean **μ** and variance-covariance matrix **K**, the expectation of *V*_*y*_ can be computed using the following general formula for quadratic forms (Searle et al. 1992):

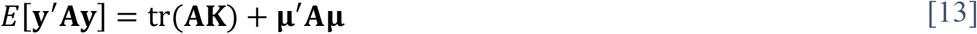

The “tr” in Eq. 13 stands for trace, which equals the sum of the diagonal elements. It follows that

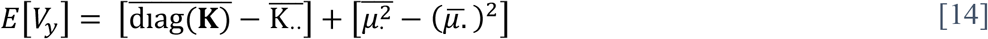

where 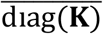 is the mean of the diagonal elements of **K**. Eq. 14 follows the convention of using an overbar to indicate averaging with respect to dotted subscripts.

For effects indexed only by genotype, such as *g*_*i*_, Eq. 14 needs to be modified to accommodate unbalanced experiments. If **x**∼MVN(**μ**, **K**), and **Z** is the incidence matrix relating **x** to the *gE* basis of the Stage 2 model, then **y** = **Zx** is the random vector for which we need to compute the expected variance. The result is identical to Eq. 14 provided the averages are interpreted as weighted averages:

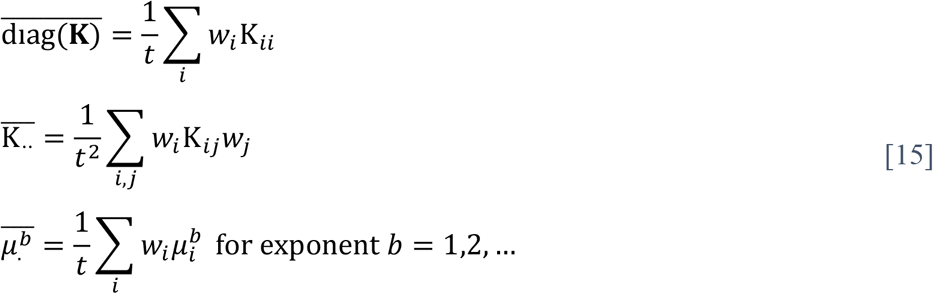

The weights *w*_*i*_ in Eq. 15 come from 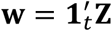 and represent the number of environments for genotype *i*.

For the multi-location model, the genotype within location variance is computed using **K** = **G⨂Γ** and weights equal to the number of times each *gL* combination is present. For a balanced experiment with *n* individuals and *s* locations, the result is

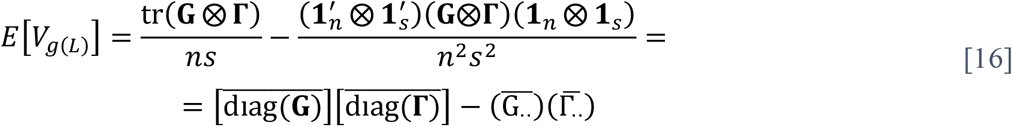

Following Rogers et al. (2021), Eq. 16 is partitioned into a main effect *V*_*g*_ plus genotype x loc interaction *V*_*gL*_. The main effect is based on the average of the 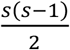 off-diagonal elements of **Γ**:

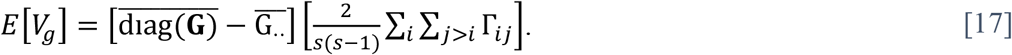

Eq. 17 is extended to the unbalanced case by using weighted averages for **G**.

### BLUP and Selection Response

Empirical BLUPs are calculated conditional on the variance components estimated in Stage 2. All Stage 2 models described above can be written in the following standard form:

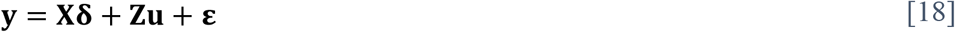

where **δ** is a vector of fixed effects (for environments, markers, and inbreeding), **u** is a vector of multivariate normal genetic effects, and **ε** is the “residual” vector (for the g x env and Stage 1 error effects). Let 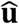 denote BLUP[**u**], which is calculated one of two ways for numerical efficiency. If the length of **y** exceeds the length of **u**, then 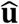 is calculating by inverting the coefficient matrix of the mixed model equations (MME; Henderson 1975). Otherwise, 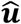 is calculated by inverting **V** = *var*(**y**) and using the following result (Searle et al. 1992):

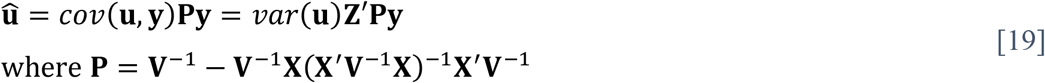

Genetic merit is a linear combination of random and fixed effects. For random effects, the structure of **u** is trait nested within individual, nested within additive vs. non-additive values. For fixed effects (ignoring the environment effects), **δ** contains trait nested within marker effects, followed by trait nested within the regression coefficient for heterosis. If **W** represents the centered matrix of allele dosages for the fixed effect markers (*n* individuals x *m* markers), **F** is the vector of genomic inbreeding coefficients, and **c** is the vector of economic weights for multiple traits or locations, then the genetic merit vector for the population is

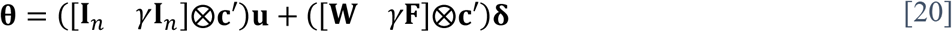

The value of *γ* depends on which genetic value is predicted: 0 for additive value, 1 for total value, and 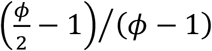 for breeding value and ploidy *ϕ* (Gallais 2003). Because BLUP is a linear operator, 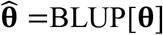 (i.e., the selection index) is given by Eq. 20 with **u** and **δ** replaced by their predicted values.

Index coefficients entered by the user are interpreted as relative weights for standardized traits (or locations). To generate the vector **c**, the software divides the user-supplied weights by the standard deviations of the breeding values (estimated in Stage 2); it also applies an overall scaling such that ‖**c**‖ = 1, which ensures predictions are commensurate with the original trait scale in multi-location models.

The reliability 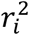 of the predicted merit 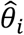 for individual *i* is the squared correlation with its true value *θ*_*i*_, which depends only on the random effects. If **u**_*i*_ represents the vector of random genetic effects for individual *i*, and **λ** denotes [1 *γ*]′**⨂c**, then the random effects component of *θ*_*i*_ is **λ**′**u**_*i*_, and the reliability is

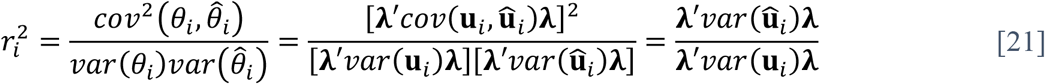

The final equality in Eq. 21 relies on the following property of BLUP: 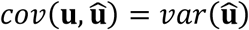. For the MME solution method, the *var*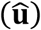 matrix is computed as *var*(**u**) − **C**_22_, where **C**_22_ is from the partitioned inverse coefficient matrix (Henderson 1975). For the **V** inversion method, 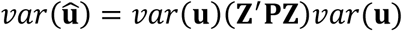 (Searle et al. 1992).

The breeder’s equation provides the expected response to truncation selection on predicted merit 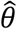. If **b** denotes the multi-trait vector of breeding values for an individual, then its predicted merit is 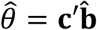 (see Eq. 20), and the multi-trait response **x** under selection intensity *i* is

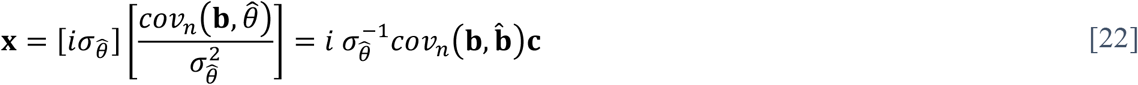

(To connect Eq. 22 with a familiar form of the breeder’s equation, the first bracketed term is the selection differential, and the second bracketed term represents heritability.) The subscript *n* on *cov*_*n*_ indicates it is the covariance with respect to the *n* individuals in the population, which differs slightly from the covariance of a vector with respect to its MVN distribution (see Appendix). As mentioned earlier, under BLUP, the latter covariance satisfies 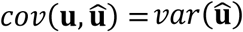. Combining this result with Appendix Eq. A5, it follows that 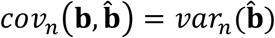, which is denoted **B**. The formula for traits *j* and *k* is

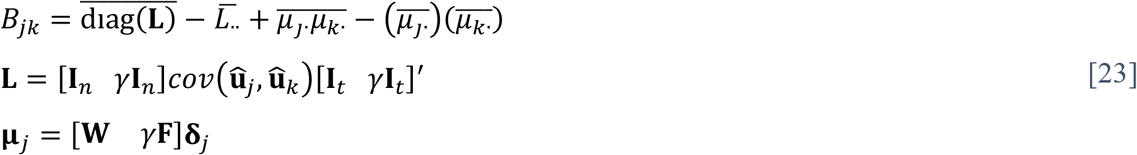

The vector 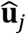 is a 2*n* x 1 stacked vector of the predicted additive and non-additive values for a population of size *n*. The calculation of 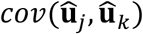 follows the same procedure described above (see Eq. 19), and the contribution from **δ** is calculated using the fixed effect estimates. Since the overall scaling of the index coefficients is arbitrary, we can impose 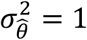. Inverting Eq. 22 under this constraint leads to an expression for the index coefficients:

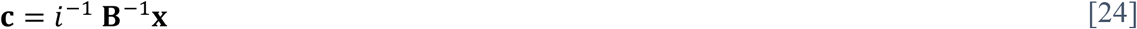

Substituting this result into 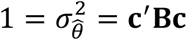 leads to an implicit equation for the response:

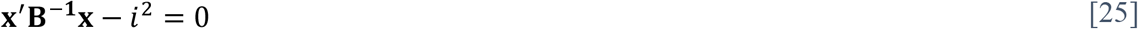

Eq. 25 is the matrix representation of an ellipsoid in *t* dimensions, which is used by StageWise to provide a geometric visualization of selection tradeoffs. (The software DESIRE [Kinghorn 2013] is an earlier example of plotting the elliptical multi-trait response.) If the response is expressed in units of genetic standard deviation, a diagonal matrix **Δ** with elements 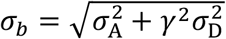 is used to rescale the matrix of the quadratic form as **ΔB**^−**1**^**Δ**. The principal axes of the ellipse are given by the eigenvectors of this matrix, and the lengths of the semi-axes equal the inverse square-root of the eigenvalues.

This geometric model provides a convenient method for implementing a restricted selection index, in which the response for some traits is constrained to be zero (Kempthorne and Nordskog 1959). From above, the change in genetic merit associated with response **x** is **c**′**x**, which is the projection of **x** onto **c** times the magnitude of **c**. For the unrestricted index, the response that maximizes genetic gain is therefore the solution of the following convex optimization problem:

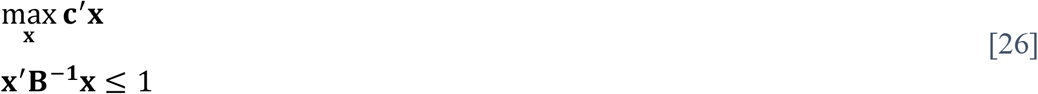

The linear inequality constraint in Eq. 26, which is convex, replaces the linear equality constraint of Eq. 25, which is not convex. This substitution is valid because the linear objective ensures the optimum is on the boundary (Boyd and Vandenberghe 2004). For the restricted index, the restricted traits are not included in the objective **c**^′^**x**, and equality or inequality constraints on the genetic gain *x*_*i*_ for restricted trait *i* are added to Eq. 26. Convex optimization is performed using CVXR (Fu et al. 2020), and the index coefficients are computed from the optimal **x** via Eq. 25 with intensity *i* = 1.

### Marker effects and GWAS

Marker effects and GWAS scores are also calculated by BLUP. Let **α** represent the *mt* x 1 vector of additive (substitution) effects for *t* traits/locations nested within *m* markers, with variance-covariance matrix **I**_*m*_**⨂Γ**(*ϕ* ∑_*k*_ *p*_*k*_*q*_*k*_)^−1^ for ploidy *ϕ* (Endelman et al. 2018). From the linearity of BLUP, the predicted multi-trait index of marker effects is 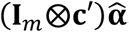, and from Eq. 19, 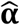 can be written in terms of the predicted additive values **â** as

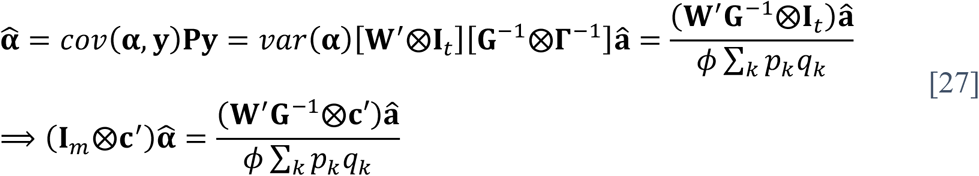

The **W** matrix in Eq. 27 is the centered matrix of allele dosages (individuals x markers). A similar result holds for relating the multi-trait index of digenic substitution effects **β** to the predicted dominance values 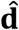 (Eq. 7):

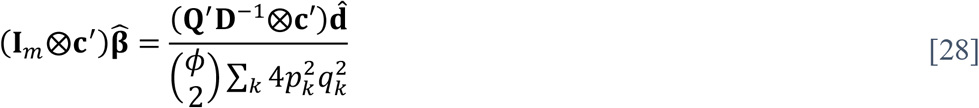

The fixed effect for inbreeding is included in 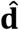 and therefore represented in the predicted marker effects.

GWAS p-values are computed from the standardized BLUPs of the marker effects, which are asymptotically standard normal (Gualdrón Duarte et al. 2014). If **w**_*k*_ denotes the *k*^th^ column of the **W** matrix, then the standard error of the predicted additive effect for marker *k* is

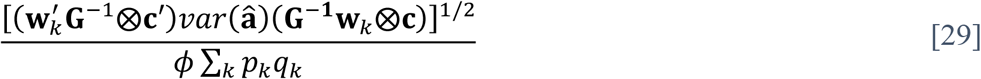

The formula for dominance effects is analogous, based on Eq. 28. StageWise provides the option to parallelize this computation across multiple cores. To control for multiple testing, the desired significance level specified by the user is divided by the effective number of markers (Moskvina and Schmidt 2008) to set the p-value discovery threshold.

### Potato data analysis

The potato dataset is an updated version of the data from Endelman et al. (2018), which spanned 2012–2017 at one location (Hancock, WI) and contained 571 clones from both preliminary and advanced yield trials. The current version spans 2015–2020 and contains 943 clones. Fixed effects for block or trial, as well as stand count, were used in Stage 1. Three traits were analyzed: total yield (Mg ha^-1^), vine maturity (1 [early] to 9 [late] visual scale at 100 days after planting), and potato chip fry color (Hunter *L*) after 6 months of storage. The G matrix was used for multi-trait analysis, instead of H, due to convergence problems with the latter.

Marker data files contain the estimated allele dosage (0–4) from genotyping with potato SNP array v2 or v3 (which contains most of v2) (Felcher et al. 2012; Vos et al. 2015). Genotype calls were made with R package fitPoly (Zych et al. 2019). Data from the two array versions were combined with the command *merge_impute* from R package polyBreedR (https://github.com/jendelman/polyBreedR). This command performs one iteration of the EM algorithm described in Poland et al. (2012) (only one iteration is needed for complete datasets at low and high density), followed by shift and scaling (if necessary) to ensure all data are in the interval [0, ploidy].

## RESULTS

The workflow to analyze data with StageWise is illustrated in Figure 1. Any software can be used to compute genotype BLUEs and their variance-covariance matrix in Stage 1. For convenience, the package has a command named *Stage1*, which can accommodate any number of fixed or i.i.d. random covariates, as well as spatial analysis using SpATS (Rodríguez-Álvarez et al. 2018). To partition genetic value into additive and non-additive components, genome-wide marker data is processed with the command *read_geno*, and the output is then included in the call to *Stage2*. After estimating the variance components with *Stage2*, the *blup_prep* command inverts either the coefficient matrix of the mixed model equations or the variance-covariance matrix of the Stage 2 response variable, whichever is smaller. This allows for rapid, iterative use of the *blup* command to obtain different types of predictions and standard errors, which are used in the calculation of reliability (i.e., squared correlation between true and predicted value) for individuals and GWAS scores for markers. Three vignettes, or tutorials, come with the software to give detailed examples of using the commands. The following results represent a condensed version of this information.

**Figure 1.**
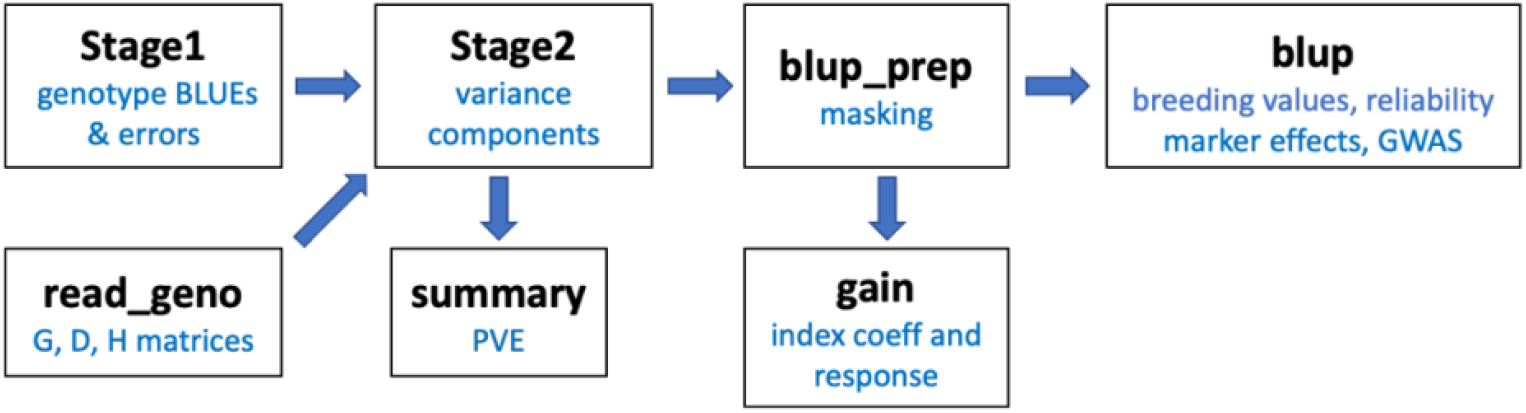
Overview of the commands and workflow in R/StageWise.

The primary dataset comes from six years of potato yield trials at a single location and includes 943 genotyped clones. The genotypic values of heterozygous clones have both additive and non-additive components. Non-additive values can be modeled in StageWise either as genetic residuals (no covariance) or as dominance values. In the context of genomic prediction, directional dominance models use inbreeding coefficients to estimate heterosis. Figure 2 compares three types of inbreeding coefficients for this population: (1) F_D_, from the directional dominance model, (2) F_G_, from the diagonal elements of the additive genomic relationship matrix, and (3) F_A_, from the diagonal elements of the pedigree relationship matrix. The F_G_ and F_D_ coefficients from the genomic models were highly correlated (*r* = 0.98) and have the same population mean, −0.08, which indicates a slight excess of heterozygosity compared to panmictic (Hardy-Weinberg) equilibrium. Although there was some concordance between the genomic and pedigree coefficients for the most inbred individuals, there was little agreement at small values of F_A_ (Fig. 2).

**Figure 2.**
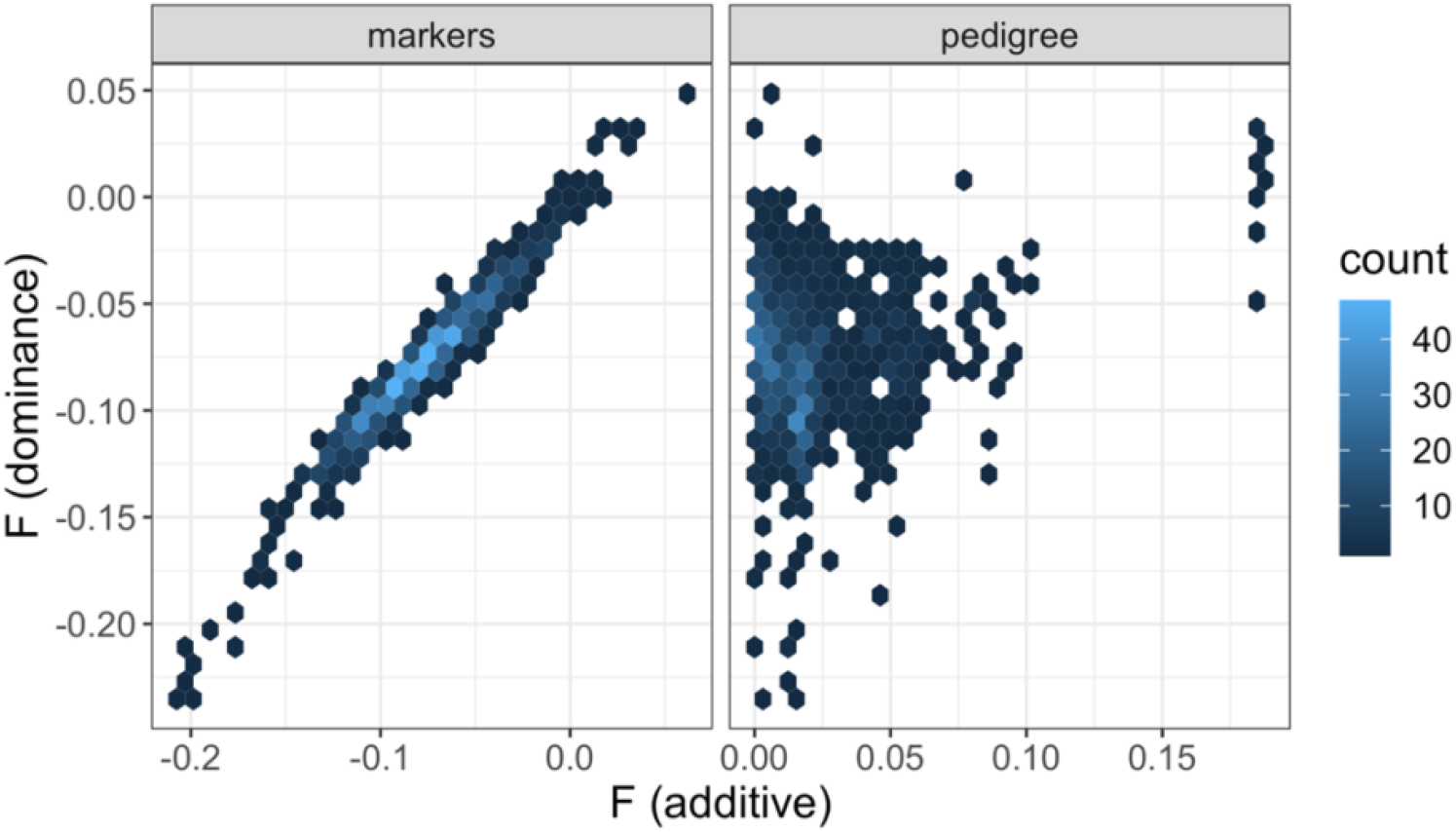
Comparison of inbreeding coefficients (F) for a population of 943 potato breeding lines. The vertical axis is computed from the dominance coefficients, and the horizontal axis is computed from the additive relationship matrix.

### Single trait analysis

Initially, the three traits in the potato dataset—total yield, chip fry color, and vine maturity— were analyzed independently. In Stage 1, broad-sense heritability on a plot basis was highest for yield (0.70 – 0.83), with similar results for fry color (0.25 – 0.74) and maturity (0.38 – 0.74) (Figure S1). The benefit of including Stage 1 errors in the Stage 2 model was assessed based on the change in AIC, which ranged from −29 for maturity to −104 for fry color (Table 1). Applying the *summary* command to the output from *Stage2* generates a table with the proportion of variation explained (PVE). The PVE for additive effects, which can be called genomic heritability, ranged from 0.34 (yield) to 0.43 (maturity) (Table 2). The PVE for dominance effects has two parts: one due to the variance of the dominance effects (“Dominance” in Table 2), and the other from variation in the genomic inbreeding coefficient (“Heterosis” in Table 2). Of the three traits, yield had the largest influence of dominance, with a combined PVE of 0.15.

**Table 1.** Akaike Information Criterion (AIC) for the Stage 2 model with vs. without inclusion of the Stage 1 errors.

|  | Yield | Fry Color | Vine Maturity |
| --- | --- | --- | --- |
| Without | 7007 | 4662 | 2018 |
| With | 6940 | 4558 | 1989 |
| Change | -67 | -104 | -29 |

**Table 2.** Proportion of variation explained for the multi-year potato dataset.

|  | Yield | Fry Color | Vine Maturity |
| --- | --- | --- | --- |
| Additive | 0.34 | 0.38 | 0.43 |
| Dominance | 0.12 | 0.04 | 0.02 |
| Heterosis | 0.03 | 0.00 | 0.00 |
| Genotype x Year | 0.30 | 0.24 | 0.16 |
| Stage 1 Error | 0.21 | 0.34 | 0.39 |

Both “Dominance” and “Heterosis” come from the directional dominance model.

StageWise has the ability for genomic prediction with the H matrix, which is a weighted average of G and A that was originally developed to use ungenotyped individuals in the training population (Legarra et al. 2009; Christensen and Lund 2010). Even when all individuals are genotyped, H may still outperform G due to the sparsity of A (Figure 3). For the potato dataset, the change in AIC with H ranged from −6 (fry color) to −13 (yield). The optimum weight for A was 0.3 for vine maturity and fry color and 0.5 for yield. As the weight for A increased, the estimate for genomic heritability (solid line in Fig. 3) also increased, at the expense of dominance (dashed line).

**Figure 3.**
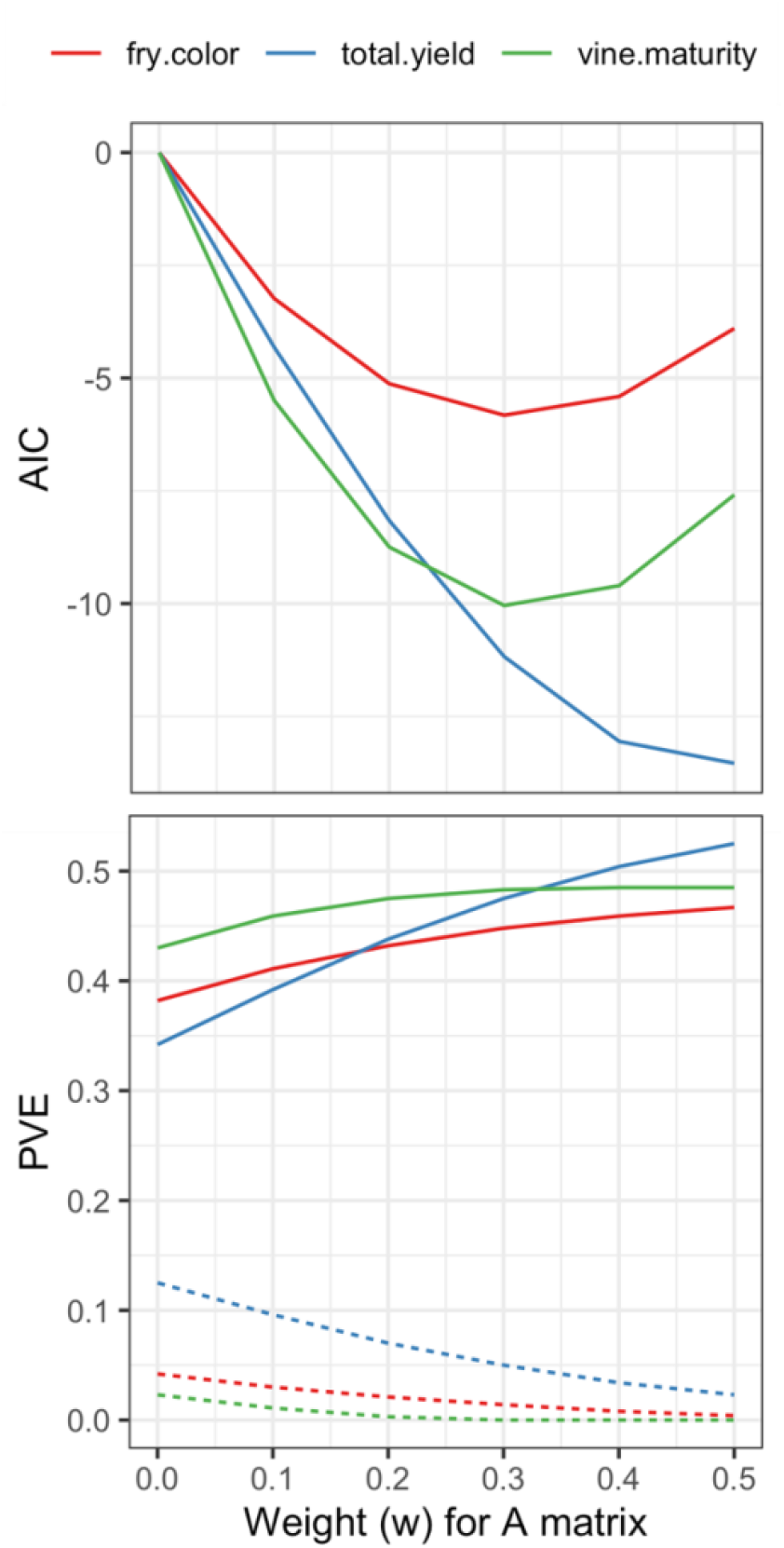
Minimizing the Akaike Information Criterion (AIC) to select the optimal weighting of pedigree (A) and marker (G) additive relationship matrices: **H** = w**A** + (1–w)**G**. The optimal weight varied by trait in a potato dataset of 943 clones. The proportion of variation explained (PVE) by the additive effects (solid line) increased with w, while the PVE for the dominance effects (dashed line) decreased.

The *blup_prep* command has an option to mask Stage 1 BLUEs, which can be used to estimate the accuracy of predicting new individuals or new environments. Figure 4 compares the reliability of genome-wide marker-assisted selection (MAS) vs. marker-based selection (MBS) for the last breeding cohort in the potato dataset. The distinction between MAS and MBS is that the selection candidates are part of the training set with MAS but not with MBS (Bernardo 2010). The reliability of 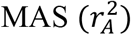 was 0.14–0.21 higher than 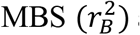 across traits. From index theory (Lande and Thompson 1990; Riedelsheimer and Melchinger 2013), the two quantities are related by

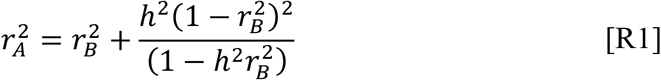

**Figure 4.**
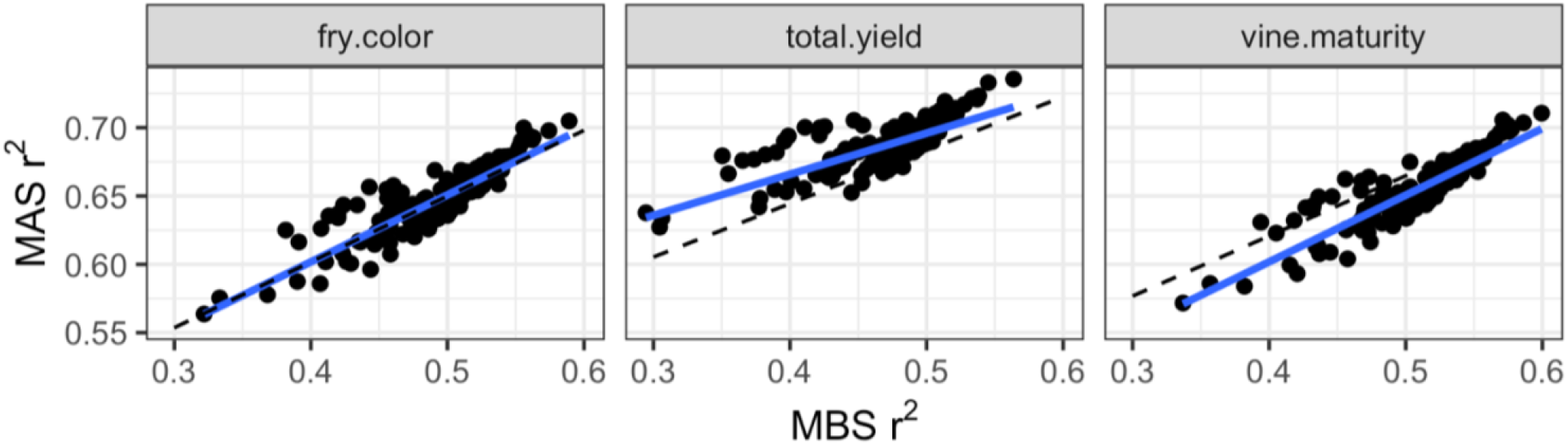
Comparing the reliability (r^2^) of marker-assisted (MAS) vs. marker-based (MBS) genomic selection in the potato dataset. Each point represents a clone from one breeding cohort, and the blue line is a linear trendline. The increased accuracy from having phenotypes for the selection candidates (MAS) was closely predicted by selection index theory (dashed line).

When used with the genomic heritability estimates from *Stage2*, this formula closely matched the data for all three traits (Fig. 4).

Although GWAS is not the emphasis of StageWise, the software can perform a fully efficient, two-stage GWAS. For the potato dataset, there was a major QTL for vine maturity on chr05 (Figure S2), in the vicinity of the well-known regulator of potato maturity *StCDF1* (Kloosterman et al. 2013). *Stage2* has an optional argument to include markers as fixed effects for major QTL. In this case, the PVE for the marker was 0.10, which represents 21% of the total additive variance.

### Multi-trait analysis

Multi-trait analysis follows the same general workflow as a single trait. In addition to the PVE, the *summary* command returns the additive correlation matrix for the traits. For the potato dataset, late maturity was correlated with higher yield (*r* = 0.57) and slightly with lighter fry color (*r* = 0.23). There was no genetic correlation (*r* = 0.00) between yield and fry color.

The “index.coeff” argument for *blup* is used to specify the selection index coefficients, which determine the relative weights of the traits (after standardization to unit variance) for genetic merit. (Because StageWise uses a multi-trait BLUP, the optimal index coefficients equal the coefficients of genetic merit.) For the potato chip market, it is reasonable to give equal weight to yield and fry color. However, naïve selection on these traits alone will generate offspring with later maturity, which is undesirable. One way to avoid this is by using vine maturity as a covariate in the analysis.

Alternatively, the *gain* command in StageWise can be used to compute the coefficients of a restricted selection index, in which the response for some traits is constrained to be zero (Kempthorne and Nordskog 1959). For a given selection intensity and *t* traits, the set of all possible responses is a *t*-dimensional ellipsoid, and *gain* shows 2D slices of it. Figure 5 shows the breeding value response for yield and maturity, as well as two line segments. The dashed red line is the projection of the index vector, and the solid blue line is the projection of the optimal response. The restricted index requires negative weight for maturity to produce zero response, which reduces the yield response compared to the unrestricted index by 0.23*iσ* (*i* is selection intensity and *σ* is the genetic standard deviation of the breeding values; Table 3).

**Figure 5.**
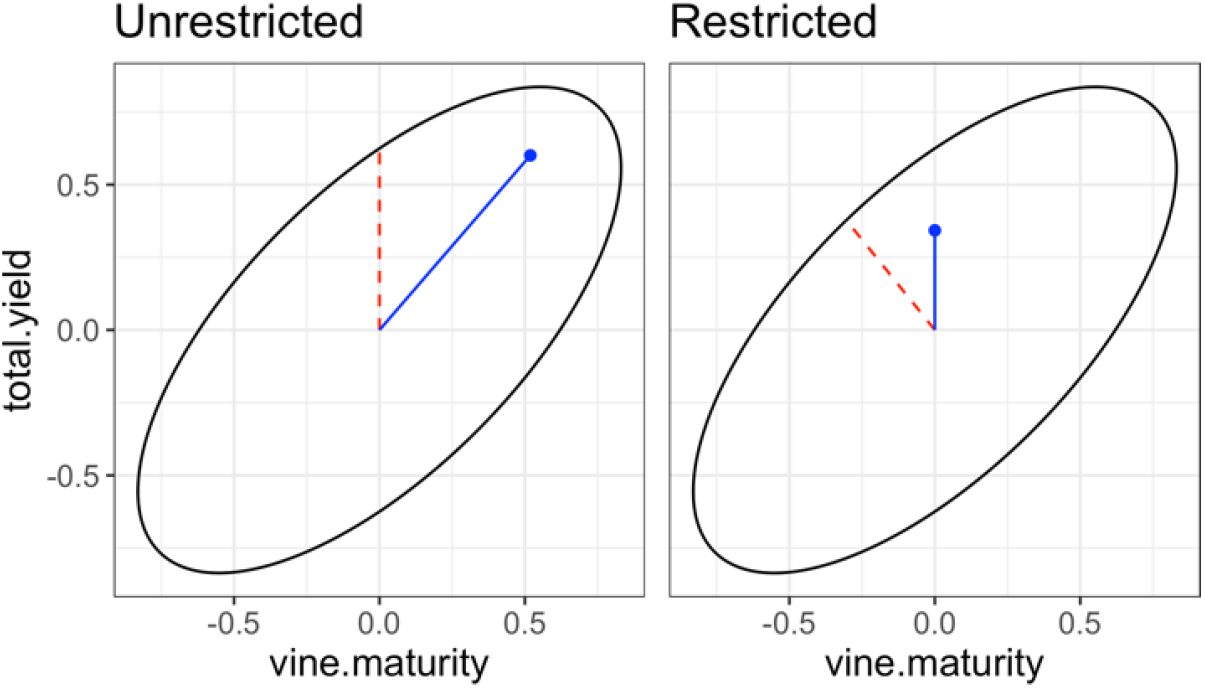
Selection response tradeoffs for yield and maturity in the potato dataset. The dashed red line segment is the projection of the index vector, and the solid blue line segment is the projection of the optimal response.

**Table 3.** Multi-trait response for potato under truncation selection, assuming yield and fry color contribute equally to genetic merit.

| Trait | Unrestricted Index |  | Restricted Index |  |
| --- | --- | --- | --- | --- |
|  | Coefficients | Response | Coefficients | Response |
| Total Yield | 0.707 | 0.53 | 0.601 | 0.30 |
| Fry Color | 0.707 | 0.52 | 0.601 | 0.51 |
| Vine Maturity | 0.000 | 0.47 | -0.527 | 0.00 |

Index coefficients are for standardized traits and scaled to have unit norm. Response is for intensity $i = 1$ , in units of genetic standard deviation.

## DISCUSSION

StageWise was designed to enhance the use of genomic prediction in plant breeding, but there are notable limitations. At present, each phenotype is associated with a single genotype identifier, which is inadequate for hybrid prediction. *Stage2* uses ASReml-R (Butler et al. 2018) for variance component estimation, which requires a paid license, but other options may become available in the future, such as sommer (Covarrubias-Pazaran 2018). The options for modeling GxE are somewhat limited, particularly for multiple traits, which assumes a uniform genetic correlation between environments.

For single trait analysis, a more complex GxE model is possible to allow for heterogenous genetic correlation between locations. The genetic covariance between locations is based on a second order factor-analytic (FA2) model (Smith et al. 2001), which offers enough statistical complexity for many applications. To assess model adequacy, the factor loadings returned by *Stage2* can visualized with the command *uniplot*, which generates a circular plot in which the squared radius for each location equals the proportion of genetic variance explained by the latent factors (Cullis et al. 2010). This functionality is illustrated in Vignette 2 using national trial data for potato (Schmitz Carley et al. 2019). At present, StageWise does not have functionality for genomic prediction with environmental covariates.

Previous research on genomic prediction in polyploids has not used directional dominance, which means inbreeding depression/heterosis was not modeled (Endelman et al. 2018; Amadeu et al. 2020; Batista et al. 2022). Heterosis explained less than 5% of the variance for yield in both potato datasets, but we should expect small PVE when there is limited variation for inbreeding. The standard deviation of F_D_ was only 0.03 for the population of 943 potato clones (Fig. 3).

From the theory of directional dominance, the average dominance coefficient is the covariate for estimating heterosis. Xiang et al. (2016) used average heterozygosity for the covariate because under a genotypic parameterization of dominance in diploids, this is equivalent to the average dominance coefficient. However, studies employing orthogonal parameterizations of dominance have also used this covariate (Aliloo et al. 2017; Yadav et al. 2021), even though heterozygosity is no longer equivalent to the dominance coefficient because the relative contribution of the genotypes to inbreeding depends on allele frequency (see Eq. 5). For example, the minor allele homozygote contributes more to inbreeding than the major allele homozygote, and the difference is *ϕ*(*ϕ* − 1)(*q* − *p*) for ploidy *ϕ* and minor allele frequency *p* = 1 − *q* at panmictic equilibrium. To give another example, simplex dosage of the minor allele in a tetraploid contributes more to inbreeding than duplex dosage only for *p* > 1/3; for *p* < 1/3, duplex dosage contributes more.

A more general approach to restricted selection indices was developed in StageWise by investigating the geometry of the problem (Eq. 26). Until now, only equality constraints have been included (i.e., specifying a certain value for genetic gain), which are amenable to solution by the method of Lagrange multipliers. StageWise uses convex optimization software to allow for both equality and inequality constraints. In many situations, inequality constraints are more appropriate than equality constraints. For example, when selecting for yield, we might accept earlier but not later maturity, which is represented by response ≤ 0. With only one constrained trait, the optimal solution corresponds to zero response, so the inequality offers no advantage. But with two or more constraints, higher genetic gains are possible with inequalities (example in Supplement).

The “mask” argument for *blup_prep* makes it easy to investigate the potential benefit of using a correlated, secondary trait to improve genomic selection. Many plant breeding programs are exploring the use of spectral measurements from high-throughput phenotyping platforms to improve selection for yield. For example, Rutkoski et al. (2016) demonstrated that aerial measurements of canopy temperature during grain fill could be used to predict wheat grain yield. Vignette 3 shows how to recreate this result in StageWise with a few lines of code.

Typically, the number of traits a breeder must consider for selection is too large to analyze jointly in *Stage2*, based on the current implementation with ASReml-R. New algorithms may alleviate this limitation in the future (Runcie et al. 2021), but in the meantime, a practical approach is to split the traits into groups for multivariate analysis based on phenotypic correlations. In the final step, multiple outputs from *blup_prep* can be combined in one call to *blup*, using an index that covers all traits (example in Vignette 3).

We should acknowledge that truncation selection on breeding value is not optimal for long-term genetic gain. The design of selection methods that conserve and exploit genetic diversity more efficiently is an exciting area of research (e.g., Toro and Varona 2010; Akdemir and Sánchez 2016; Goiffon et al. 2017). Although such methods are not currently available in StageWise, the additive and dominance marker effects returned by the software can be used to implement them.

## Declarations

## Acknowledgments

I would like to thank potato breeding colleagues across the US for contributing germplasm used in this study, Grace Christensen for assistance with genotyping, and the UW-Madison Hancock and Rhinelander Agricultural Research Stations.

## Data Availability

The potato datasets and vignettes are distributed with the StageWise software, which is available at https://github.com/jendelman/StageWise under the GNU General Public License v3.

## APPENDIX

The objective is an expression for the expected covariance between two quantities of a population of size *n*, represented by multivariate normal vectors **x**_1_∼MVN(**μ**_1_, **K**_1_) and **x**_2_∼MVN(**μ**_2_, **K**_2_), with covariance **L**:

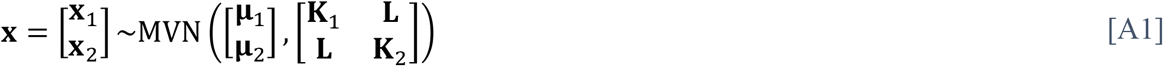

Noting that **x**_1_ = [**I**_***n***_ **0**]**x** and **x**_2_ = [**0 I**_***n***_]**x**, the population covariance is (cf. Eq. 12)

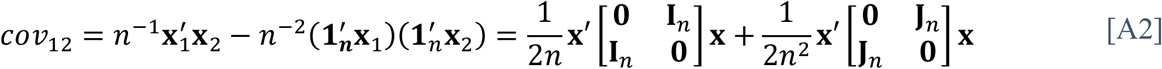

where 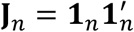 is a *n* x *n* matrix of ones. Using Eq. 13, the expectation of the first quadratic form in Eq. A2 is

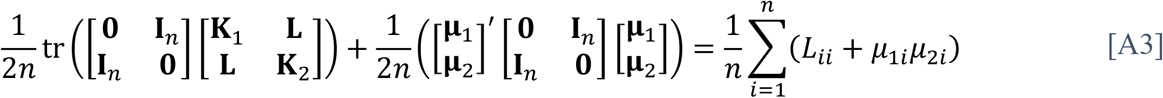

The expectation of the second quadratic form in Eq. A2 is

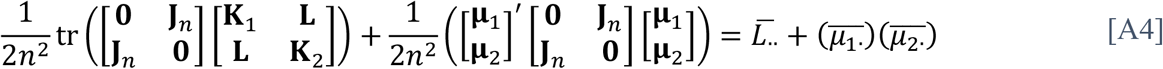

Putting Eq. A3 and A4 together, the expected covariance is

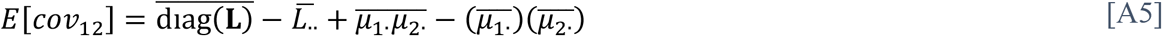

As in the Methods, for partitioning covariance on a *gE* basis, the unbalanced nature of the experiment is accounted for by computing the covariance between vectors **y**_1_ = **Zx**_1_ and **y**_2_ = **Zx**_2_, where incidence matrix **Z** maps *n* individuals to *gE* instances. If **y** denotes the stacked vector [**y**_1_ **y**_2_]′, then

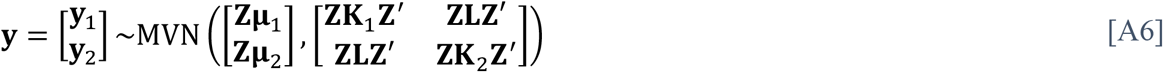

Replacing **x** with **y** in Eq. A2, the result for expected covariance follows Eq. A5 but using averages weighted by the number of environments per genotype.

## Supplementary Figures

**Figure S1.**
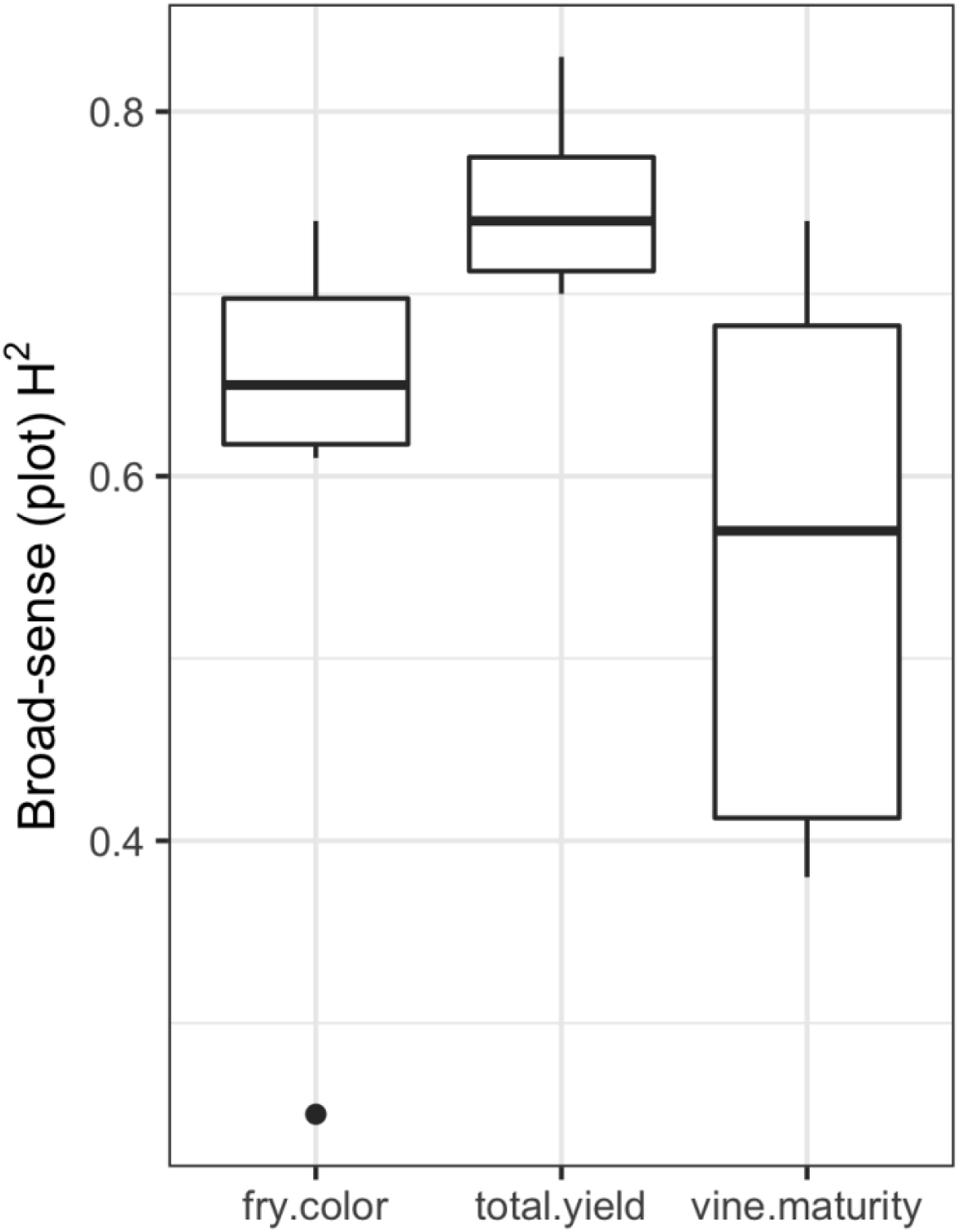
Distribution of the broad-sense heritability (plot basis) by year, from six years of potato yield trials at one location (Hancock, WI).

**Figure S2.**
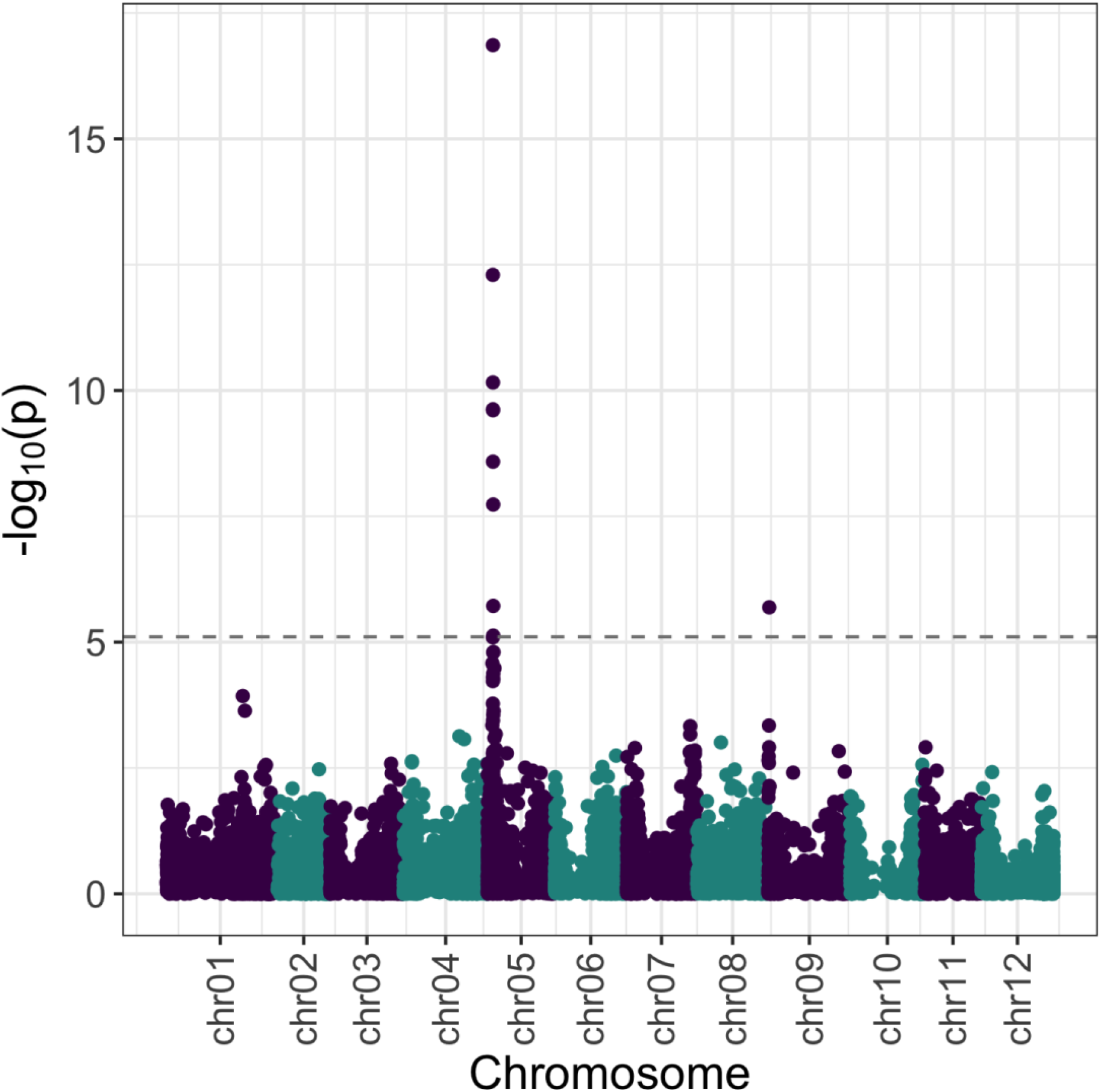
Genome-wide association results for potato vine maturity, using an additive model. The dashed line is the discovery threshold at significance level 0.05, corrected for multiple testing based on the effective number of markers.

## Supplementary File: Restricted Selection Index

StageWise uses a new approach to compute the restricted selection index coefficients based on convex optimization software, which allows for inequality and equality constraints on the response. Equality constraints have been historically used for mathematical convenience, even if an inequality better represents the situation. This applies when a response in only one direction (e.g., positive) is acceptable even though no economic weight is given to the trait.

With a single restricted trait, the optimum solution corresponds to zero response, so the inequality constraint provides no advantage. However, with two or more restricted traits, a greater response for the target traits can be observed when using inequality constraints, depending on the genetic correlations.

The gain function in StageWise can be used to illustrate this phenomenon. This command was introduced in Vignette 3, in which case the first argument was the output from the blup_prep command. The gain function computes the matrix of the quadratic form from this data (*B*^−1^ in Eq. 25, rescaled by the genetic standard deviations), but it is also possible to directly supply this matrix. For simplicity, we will consider selection on true values without dominance, in which case the matrix is the inverse of the additive correlation.

Consider selection on one focal trait, which is correlated with two other traits (values of 0.3 and 0.6) for which a positive response is undesirable. The correlation between the restricted traits will be varied from 0 to 0.9, and the response for the target trait will be computed for two different restricted indices: inequality vs. equality constraints.

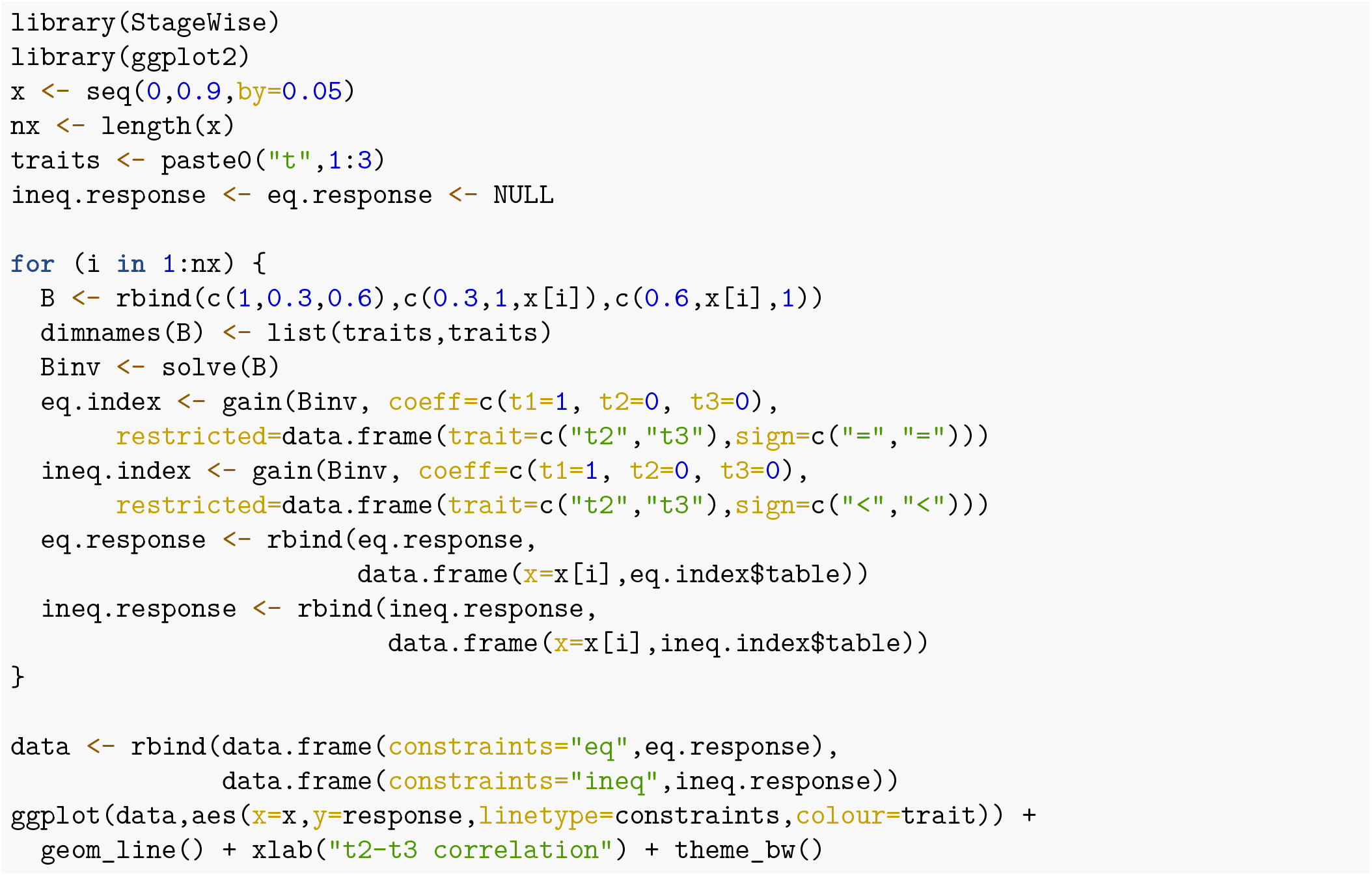

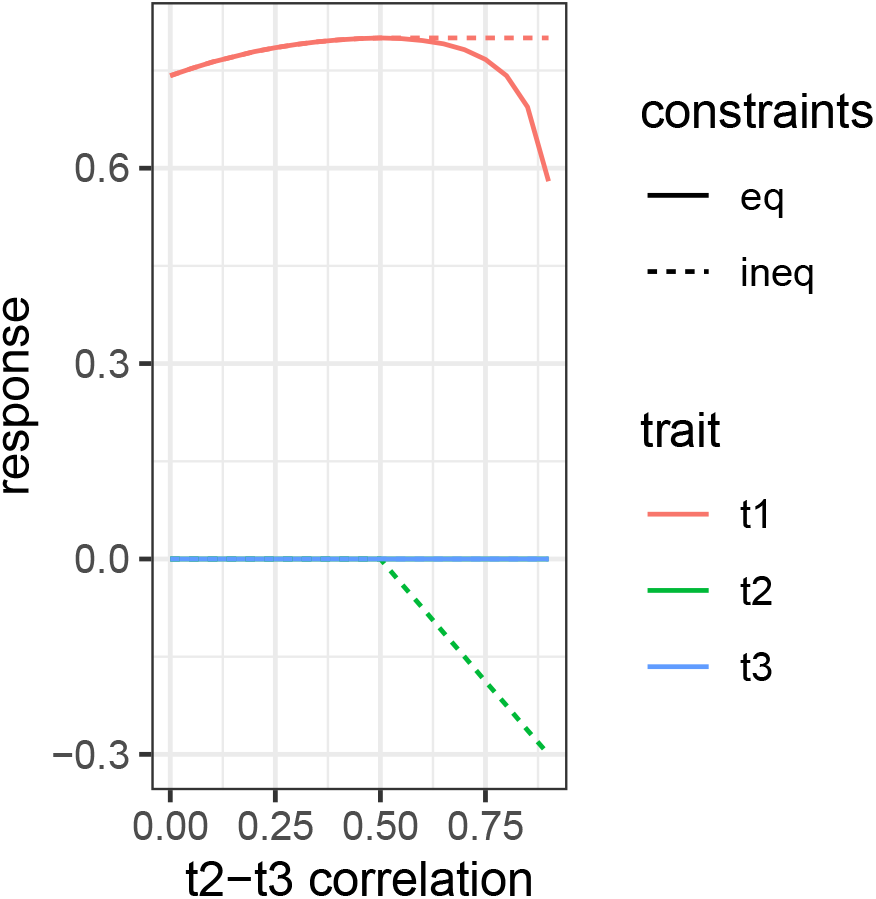

The figure shows that once the t2-t3 correlation exceeds 0.5, higher gains in t1 are possible by allowing a negative response for t2, which is the trait with the lower correlation with t1.

